# Spatial and temporal localization of *Serratia ureilytica* causing cucurbit yellow vine disease in cucurbits indicates phloem-associated colonization and systemic movement

**DOI:** 10.64898/2026.02.18.706665

**Authors:** Kensy D. Rodriguez-Herrera, Elise Boisvert, Margaret H Frank, Christine D. Smart

## Abstract

Cucurbit yellow vine disease (CYVD), caused by the bacterium *Serratia ureilytica*, is a phloem-associated disease of cucurbits. This study characterized the spatial and temporal distribution of *S. ureilytica* in *Cucurbita pepo* cultivar ‘Delicata’ plants under greenhouse conditions using a GFP-tagged isolate (P01). Seedlings were sampled weekly for four weeks. Transverse sections from the stem, petiole, leaf, shoot apex, and root were imaged by laser scanning confocal and fluorescent dissecting microscopy. In parallel, bacterial abundance in each plant tissue was assessed by quantifying colony-forming units (CFU) via droplet plating over a 4-week time course. Across plant tissues and time points, *S. ureilytica* fluorescent signal was primarily concentrated in the inner and outer periphery of the bicollateral vascular bundles, with higher magnification images revealing mainly symplastic localization within phloem-associated cells. Consistent with the imaging results, bacterial quantification data showed a high abundance of CFUs in the main stem across weeks, with an irregular pattern of presence in the distal tissues at later time points. These results suggest that *S. ureilytica* is predominantly localized within phloem-associated cells and spreads both acropetally and basipetally during infection.

## Introduction

*Serratia ureilytica* (formerly known as *Serratia marcescens*) is a relatively understudied vascular-based plant pathogen that infects cucurbit crops, causing Cucurbit Yellow Vine Disease (CYVD) (Mphande et al. 2025b; Rascoe et al. 2003). *S. ureilytica* is transmitted by the squash bug (*Anasa tristis*), a common pest of cucurbits (Bextine et al. 2001; Bonjour and Fargo 1989; Pair et al. 2004), as well as cucumber beetles (Mphande et al. 2025b). Symptoms of CYVD include yellowing, leaf scorching, stunting, and honey-brown discoloration of the phloem (Bruton et al. 2003, 1998a). CYVD was first reported in 1988 in Texas and Oklahoma and has since spread to several states in the US, including recent reports from New York and Iowa (Mphande et al., 2024a; Rodriguez-Herrera et al., 2023). Losses due to CYVD have been reported from 5%-100% (Zhang et al. 2005). Previous work proposes that *S. ureilytica* is present in phloem sieve tubes, the intercellular space of parenchyma, and xylem vessels using scanning electron micrographs and fluorescence microscopy (Bruton et al. 1998b; Luo 2006).

Bacterial colonization of plant vascular tissue systems is a well-documented route for plant pathogens. One such example is *Erwinia tracheiphila*, the causative agent of bacterial wilt in cucurbits (Rojas et al. 2015). This pathogen colonizes the xylem, where it clogs vessel elements(Rojas et al. 2015). The infection ultimately blocks water transport, causing rapid wilting and, often, death (Rojas et al. 2015). Huanglongbing (HLB), caused by ‘*Candidatus Liberibacter asiaticus*’, is another vasculature-localized disease that causes tremendous crop loss in citrus groves (Gottwald 2010; Perilla-Henao and Casteel 2016). This pathogen resides in phloem sieve tube cells, where it consumes host carbon sources, replicates, and clogs phloem transport (Ding et al. 2015). Another example of a phloem colonizer is *Candidatus Arsenophonus phytopathogenicus*, which causes low-sugar syndrome in sugar beets and marginal chlorosis in strawberries. This pathogen can metabolize sugars from phloem sap as its sole source of nutrition (Mahillon et al. 2025). Phytoplasmas are another, well-described class of pathogens that colonize host sieve elements. Phytoplasmas can adhere to cell surfaces, move passively through the phloem, and, in some cases, such as *Candidatus Phytoplasma asteris*, which causes aster yellows disease, manipulate plant development to induce distinctive phenotypes through secreted effector proteins (Bendix and Lewis 2018). The majority of phloem-colonizing pathogens share commonalities: they are found in phloem sieve tube elements and are generally unculturable in artificial media (Bendix and Lewis 2018; Haider et al. 2024; Mahillon et al. 2025). Although several examples of phloem-colonizing pathogens are known, they all infect sieve elements; little is known about pathogens that colonize other phloem-associated cells.

Vascular plants transport water and nutrients via xylem and phloem (Agustí and Blázquez 2020). Xylem transports water and solutes from the soil upward, whereas phloem transports photoassimilates and signaling molecules throughout the plant, from source to sink tissues (Taylor et al. 2009). In flowering plants, the phloem is composed of enucleate sieve elements through which phloem sap travels, companion cells that support sieve tubes in phloem sap transport, and which are extensively connected with sieve tube elements via specialized plasmodesmatal pores to form sieve element-companion cell complexes (Van Bel 2003; Haider et al. 2024; Reidel et al. 2009; De Schepper et al. 2013; Turgeon 2017). Companion cells are relatively small and contain the regulatory and metabolic mechanisms for the transport of phloem sap (Braun 2022; Perilla-Henao and Casteel 2016). Phloem parenchyma are metabolically active cells within the vascular bundle that can function as transient or long-term storage for carbohydrates and other solutes (Esau 1969; De Schepper et al. 2013).

Phloem physiology can be broken down into three functions: collection, transport, and release (de Schepper et al., 2013). In source tissues (i.e., mature leaves) where photosynthesis occurs, sugars are produced and loaded into the sieve element-companion cell complexes of collection phloem, passed via transport phloem throughout the plant, and finally offloaded into sink tissues via the release phloem. Sink tissues change throughout development and can include meristems, new leaves, and fruits, but most sugars are directed to the roots. According to the long-accepted Münch theory, this mass flow is driven by a pressure gradient, with high solute concentrations at the source and low concentrations at the sink providing the necessary force (Münch, 1930; de Schepper et al., 2013). One key modification to this theory is the “leakage retrieval mechanism,” in which loading and unloading occur dynamically between sieve element-companion cell complexes and phloem parenchyma throughout the transport phloem (Van Bel 2003; De Schepper et al. 2013; Thorpe et al. 2005). This model, which helps to explain how the pressure gradient can be tightly regulated, also provides a framework for understanding how bacteria utilizing the phloem can colonize not only terminal sink tissues, but also lateral exchange sites along the transport pathway (De Schepper et al. 2013). A notable feature of Cucurbit vasculature is its bicollateral vascular bundles, in which phloem occurs on both the internal and external sides of the xylem, rather than only externally as in a typical vascular bundle (Turgeon 2017; Zhang et al. 2012).

There is very little research on the localization of *S. ureilytica* in cucurbits. Using electron microscopy, Bruton et al. 1998 reported that, once in the plant, S*. ureilytica* moves through the phloem sieve tube elements. Recent studies have examined this bacterium in the main stem at a single time point by looking at the number of bacteria in different tissues (Mphande et al. 2025a). However, it is still unclear how the bacteria colonize the plant over time. By using fluorescence microscopy and sampling at multiple developmental stages, this study provides valuable new insights into *S. ureilytica* localization and movement as the infection spreads throughout the plant. This project had the following objectives: i) Determine the cellular and anatomical localization of *S. ureilytica* in squash plants at weekly intervals after inoculation using dissecting and confocal microscopy, and ii) quantify the distribution of *S. ureilytica* in different plant tissues throughout time using bacterial quantification methods.

## Materials and Methods

### Bacterial growth and storage

Two isolates were used for this experiment: Isolate 22212 was recovered from Ontario County, New York, at the AgriTech Research Station (Geneva, NY) in 2021, and is known to cause typical CYVD symptoms following greenhouse inoculation of plants (Rodriguez-Herrera et al. 2023). The green fluorescent protein (GFP)-tagged isolate (P01) was received from Elizabeth Little, University of Georgia, and was isolated in Georgia in 2015 (Besler 2014; Besler and Little 2017). The GFP insertion was performed in the Gerardo Lab at Emory University, as described by Mendiola et al. (2022). Isolates were stored at -80 °C in Microbank tubes (Pro-Lab Diagnostics, Ontario, Canada). To grow them from storage, three to four beads from each Microbank tube were plated onto Luria Broth (LB) agar (Bertani 1951) supplemented with tetracycline (20 μg/ml) (Mphande et al. 2025b) and incubated overnight.

### Plant cultivation

Seeds (n=100) of winter squash (*Cucurbita pepo* cv Delicata; Johnny’s Selected Seeds, Winslow, ME, USA) were germinated in deep-well 50-cell flats (T.O. Plastics, Clearwater, MN, USA) filled with soilless potting mix (Lambert’s LM-3 All Purpose Mix; Quebec, Canada). Once the seedlings had emerged (approximately 7 days after planting), they were transplanted into two-gallon pots (Grower’s Solution One Gallon Black Trade Pots; Cookeville, TN, USA). All plants were grown in Cornell University’s greenhouse facilities under a 16-hour photoperiod and day/night temperatures of 23 °C/21 °C.

### Comparison of isolates

In all experiments, isolate 22212 was included as a positive control due to its known pathogenicity on cucurbits. Isolates P01-GFP and 22212 were compared to confirm their pathogenicity and inoculation success rate.

### Inoculations

Four treatments were established: Isolate P01-GFP, isolates 22212, mock-inoculated control, and an uninoculated control. Isolates P01-GFP and 22212 were resuspended in phosphate-buffered saline (PBS) to a 1.0 optical density at 600 nm (OD600) and amended with Silwet L-77 silicone surfactant (Momentive Performance Materials, Niskayuna, NY) at a concentration of 0.02% (Mphande et al. 2024). Approximately 10 days after planting, inoculation was performed by withdrawing 300 μL of inoculum using a 26 ½ gauge needle attached to a 1 ml syringe (Becton, Dickinson & Co., New Jersey, US) and injecting it into the stem three times at a 45° angle, releasing the inoculum gently (Mphande et al. 2024). A new syringe was used for each treatment, and the needle was sterilized with 70% ethanol for 1 minute and rinsed in deionized water for 5 seconds after each plant inoculation. Each experiment was repeated once.

### Sampling

For each experiment, plants were destructively sampled at zero, seven, fourteen, twenty-one, and twenty-eight days post-treatment. Samples were collected from six locations on each plant: stem 1 cm below the soil line, 1 cm above the soil line, 2 cm above the soil line, and from the first petiole, the first leaf, and the shoot apex. Root tissue was only collected for fluorescence-based image localization. To confirm *S. ureilytica* presence in every plant sample, a direct PCR was conducted using *S. ureilytica-*specific primers (A79F-A79R) (Zhang et al. 2005).

### Bacteria localization using imaging

Samples for imaging were taken from plants inoculated with *S. ureilytica* isolate P01 (GFP), and from mock-inoculated controls. Plants were inoculated as previously described, and samples were collected for imaging at the specified time points. Transverse sections of the stem were taken from **t**he six tissue types described above, with an additional sample collected from the root, approximately 2 cm below the root-stem junction. Samples were sectioned by hand and mounted in water on a slide with a 0.17 mm coverslip (Avantor, Radnor, PA, USA). To assess the spatial distribution of *S. ureilytica* within tissue cross sections, samples were imaged using a Leica, M205 FCA fluorescence dissecting microscope equipped with an EL6000 light source and a K5 camera, with LAS X software (Leica Microsystems, Wetzlar, Germany). Exposure time and detector gain were adjusted to avoid signal saturation. Cellular localization was characterized using an LSM880 upraight Axio Examiner Z1 laser-scanning confocal microscope, with Zen image acquisition software (Zeiss, Oberkocehn, Germany). GFP fluorescence was excited using a 23 mW ArgonRemote laser and MBS 488 dichroic mirror (Zeiss, MBS-488) with an excitation peak at 488nm and an emissions peak at 545.5nm. Laser power was set to 20% of maximum output, and detector gain was set to 450, except where explicitly noted in images specifically depicting autofluorescence of plant-endogenous compounds (Supplementary Fig. 1).

To distinguish apoplastic versus symplastic compartmentalization, plants were grown and inoculated with *S. ureilytica* isolate P01 (GFP) or mock-inoculated as described above. One sample type (1 cm below the soil line) was selected because it had consistently shown a distinct GFP signal across all time points. Samples were harvested 21-days post inoculation and stained with Calcofluor White (Biotium, Fremont, CA, USA; 5mM in water), a fluorescent dye that binds to β-glucans in cell walls, for visual cell segmentation. Samples were hand-sectioned as described above, then incubated with the dye for 30 seconds, and rinsed with deionized water immediately before imaging. Samples stained with Calcofluor White were simultaneously imaged using two channels: the GFP channel (described above, but with laser power reduced to 19%), and a Calcofluor White-specific channel, using a 405-nm diode laser at 30% power, with an excitation peak at 405 nm and emission peak at 467 nm. Detector gain was adjusted at different magnifications (10x, 20x, and 63x) to optimize signal-to-noise ratios and avoid saturation. Brightfield images were also collected for all samples.

Image processing for all confocal and dissecting-scope images was performed in Fiji (ImageJ v. 1.54; Schindelin et al. 2012). Image stacks were imported, and channels (GFP, Calcofluor White, and brightfield) were merged using the ‘Merge Channels’ function for visualization and ROI identification.

### Quantification of *S. ureilytica*

Three plants for each of the four treatments were destructively sampled at the same time points and sampling locations, with exception for root samples, which were excluded from CFU quantification. Plant tissue samples (0.5 g) were surface-sterilized with 70% ethanol for 1 minute, followed by a 20-second rinse with sterile distilled water. The sterilized tissue was macerated in 500 µl of sterile distilled water. A serial dilution series was performed using the steps described by (Herigstad et al. 2001; Slack et al. 2021) resulting in five 10-fold serial dilutions. The droplet plating method was employed using LB agar plates supplemented with tetracycline (20 μg/ml), (Mphande et al. 2025b). For each dilution, 10 drops each with 10 μL were distributed across two plates (5 per dilution per plate, n=30 droplets per plate). Plates were incubated at 28°C and checked for bacterial growth after 24 hours. The dilution that gave 3-30 colonies per drop was used for counts (Herigstad et al. 2001). All colonies in the 10 drops were counted (5 drops × 2 plates) at the countable dilution. Bacterial colony forming units (CFU/ml) were calculated by multiplying the total number of colonies counted on the plate by the dilution factor, then dividing the result by the volume of the plated sample.

### Data analysis for bacterial colonization

Data were analyzed using a generalized linear mixed model (PROC GLIMMIX, SAS Institute, 2013) with adaptive quadrature (colony forming units per ml data) methods (Gbur et al., 2012). A log model was used to represent the data. The best fit for the model was based on the lowest Akaike information criterion (AIC) value. All models included Plant ID, Treatment, Isolate, Sample Location, Week, and their interaction as fixed effects, and replication as a random effect. When a significant interaction effect was detected, the SLICEDIFF function was used to compare treatments. The model Kenward–Roger method to estimate denominator degrees of freedom (Kenward and Roger 1997). Mean separations were calculated using Tukey’s Honest Significant Difference [HSD] test at α = 0.05. For this analysis, if colony growth was not observed in any tissue of a plant, the plant was excluded from the data analysis.

## Results

### Comparison of isolates

Isolations were obtained from all plants inoculated for both imaging and bacterial density experiments from six plant sampling locations. *S. ureilytica* was consistently isolated from the main stem (1 cm below, 1 cm above, and 2 cm above the inoculation site), and it was also found, although less frequently, in the petiole, leaf base, and apex (**Table 1**). Isolations were successful in recovering *S. ureilytica* even when inoculated plants showed no symptoms. For example, while 22% of plants inoculated with isolate P01 (GFP-tagged) showed CYVD-like symptoms (yellowing, leaf scorching, and stunting), the isolate was recovered from 91% of inoculated plants. Likewise, 41% of the plants inoculated with the positive control (isolate 22212) showed symptoms, and the isolate was recovered from 100% of the inoculated plants. No *S. ureilytica* was recovered from isolations of the uninoculated and/or mock-inoculated plants.

**Table 1.** *Serratia ureilytica* isolations from inoculated plants with isolate P01 from one to four weeks post-inoculation in different plant sampling locations of *Cucurbita pepo* plants (var. Delicata)

| Plant Sampling<br>location | Isolations Confocal trials |  |  |  | Isolation bacteria counting trials |  |  |  | Total<br>Isolations<br>(n=45<br>plants) |
| --- | --- | --- | --- | --- | --- | --- | --- | --- | --- |
|  | 7 DPI<br>(n=4) | 14 DPI<br>(n=7) | 21 DPI<br>(n=5) | 28 DPI<br>(n=6) | 7 DPI<br>(n=5) | 14 DPI<br>(n=6) | 21 DPI<br>(n=6) | 28 DPI<br>(n=6) |  |
| 1 cm below | 3 | 6 | 4 | 6 | 5 | 5 | 4 | 6 | 39 |
| 1 cm above | 3 | 6 | 4 | 6 | 5 | 6 | 5 | 6 | 41 |
| 2 cm above | 4 | 6 | 3 | 6 | 5 | 6 | 5 | 6 | 41 |
| Petiole | 2 | 3 | 1 | 1 | 3 | 3 | 3 | 3 | 19 |
| leaf base | 1 | 1 | 1 | 1 | 0 | 0 | 2 | 2 | 8 |
| apex | 1 | 1 | 1 | 1 | 0 | 0 | 2 | 3 | 9 |

### Bacterial localization using imaging

Transverse sections from the seven established locations were destructively sampled from inoculated and control cucurbit plants (var. *Delicata*) each week for 28 days. In inoculated plants, *S. ureilytica* localization was consistently observed in phloem-associated cells across the main stem and in distal tissues (**Table 1**). Dissecting scope images showed that *S. ureilytica* P01-GFP was concentrated in both the interior and exterior regions of each vascular bundle, consistent with the bicollateral anatomy of phloem tissue in cucurbits (**Fig. 2**). GFP signal was detectable to some degree in all sample locations across time points. Notably, the signal at 1 cm below soil level and in the root was consistently stronger than in distal shoot samples (**Fig. 2**). In these belowground sample locations the tagged *S. ureilytica* inoculate appeared to colonize a larger proportion of the host cells at three and four weeks post-inoculation relative to earlier timepoints **(Fig. 2**). Due to the destructive nature of sampling, the dynamics of disease progression could not be tracked in individuals over time. Fluorescence in control or mock-inoculated plants was only observed around the cell walls of vessel elements, which have known autofluorescent properties (Donaldson 2020) (**Supplementary Fig. 1**). This autofluorescence was clearly distinguishable from GFP signal based on its localization and faint emission (**Supplementary Fig. 1**).

**Fig 1.**
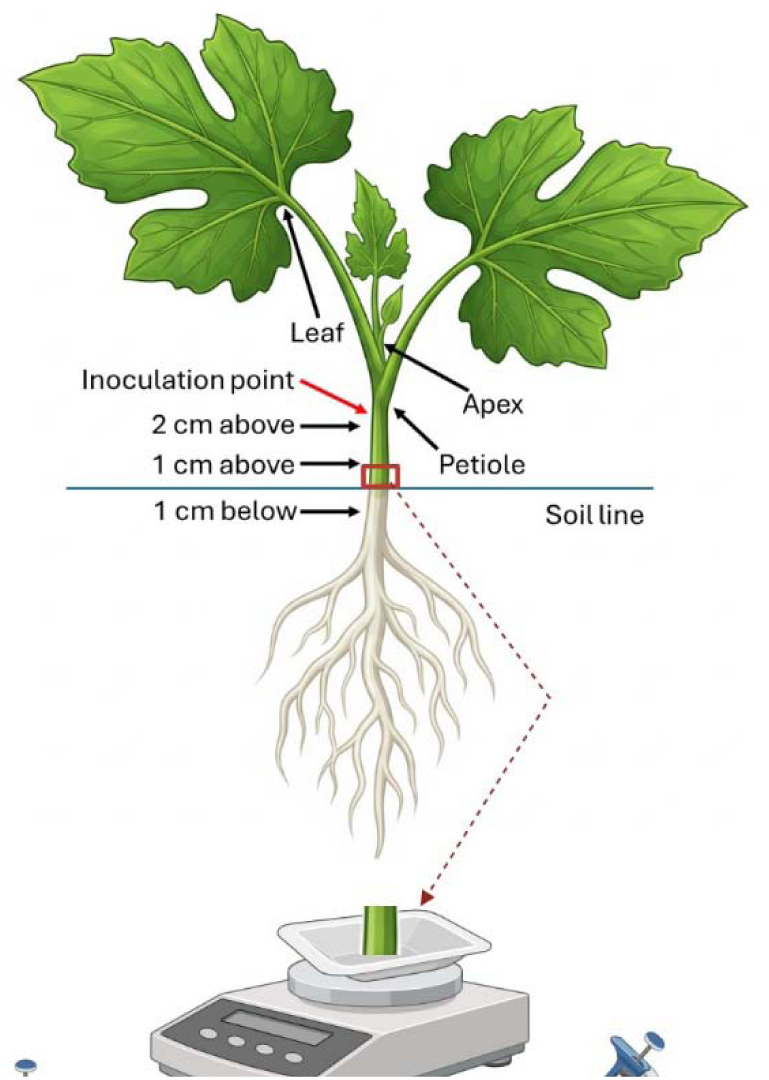
Description of methodology for bacteria quantification using the dilution plating method. Created in https://BioRender.com

**Fig 2.**
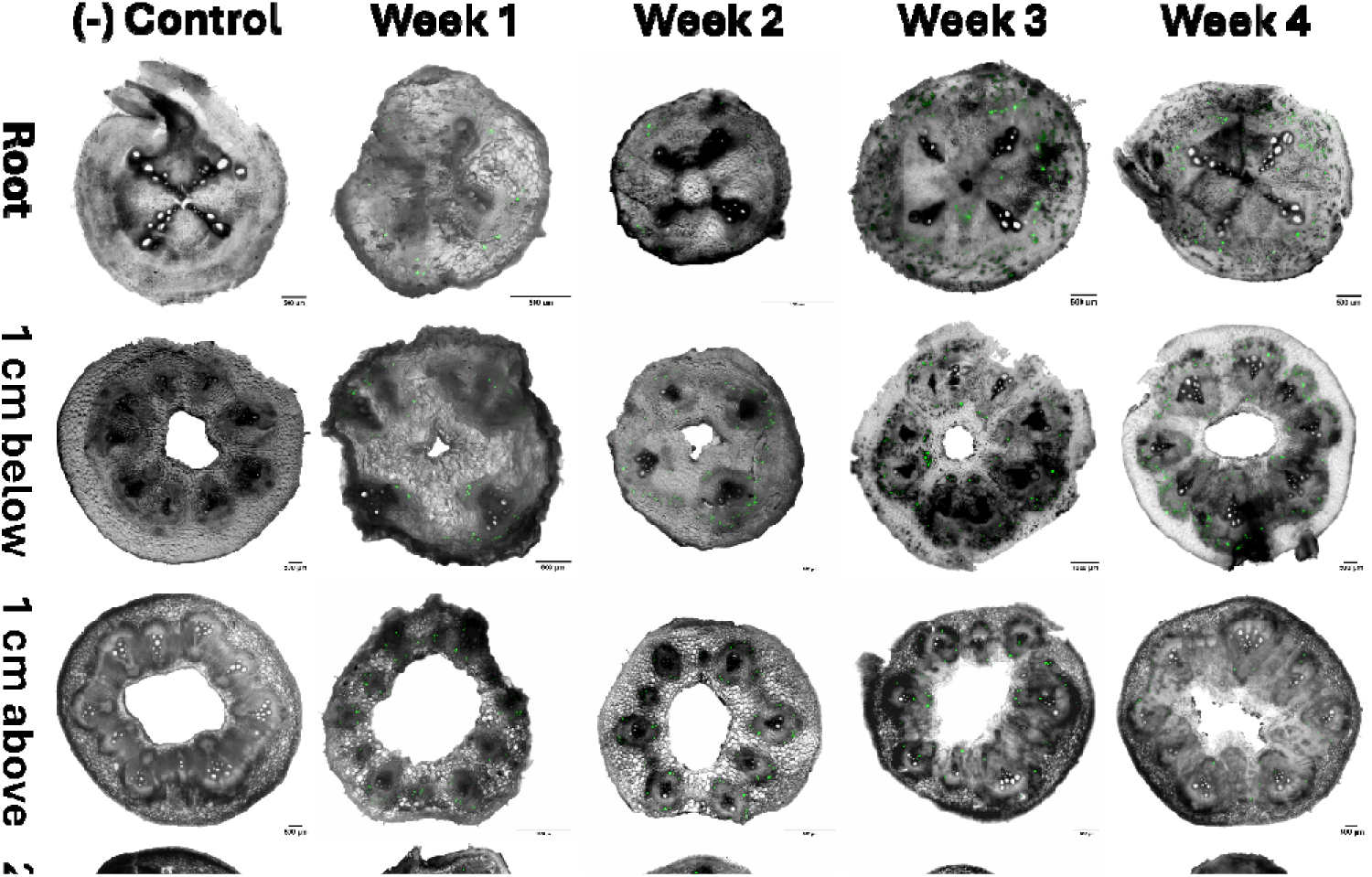
Dissecting scope images showing the localization of the green fluorescence protein (GFP)-tagged *Serratia ureilytica* in *Cucurbita pepo* plants. In six tissue types across 4 weeks. Plants were destructively sampled. Scale bar 1000 μm.

Dissecting scope imaging enabled visualization of *S. ureilytica* colonization in entire transverse sections, whereas confocal microscopy resolved signal localization at the cellular level. Confocal images taken at 10x magnification confirmed *S. ureilytica* localization within the vascular bundles and provided additional context that the signal localized to a subset of cells, scattered throughout phloem tissue, rather than colonizing in aggregated host cell clusters **(Fig. 3**). Images taken at 10x, 20x, and 63x of the lower stem (1 cm below the soil line) and stained with Calcofluor White to distinguish cell walls, revealed that the *S. ureilytica* GFP signal was predominantly symplastic, with some signal putatively localized to apoplastic regions (**Fig. 4**). Colonized cells were frequently irregular in shape and typically smaller than surrounding cells. Brightfield images also showed that colonized cells along with neighboring uncolonized cells have a darker, shadow appearance **(Fig. 4).**

**Fig. 3.**
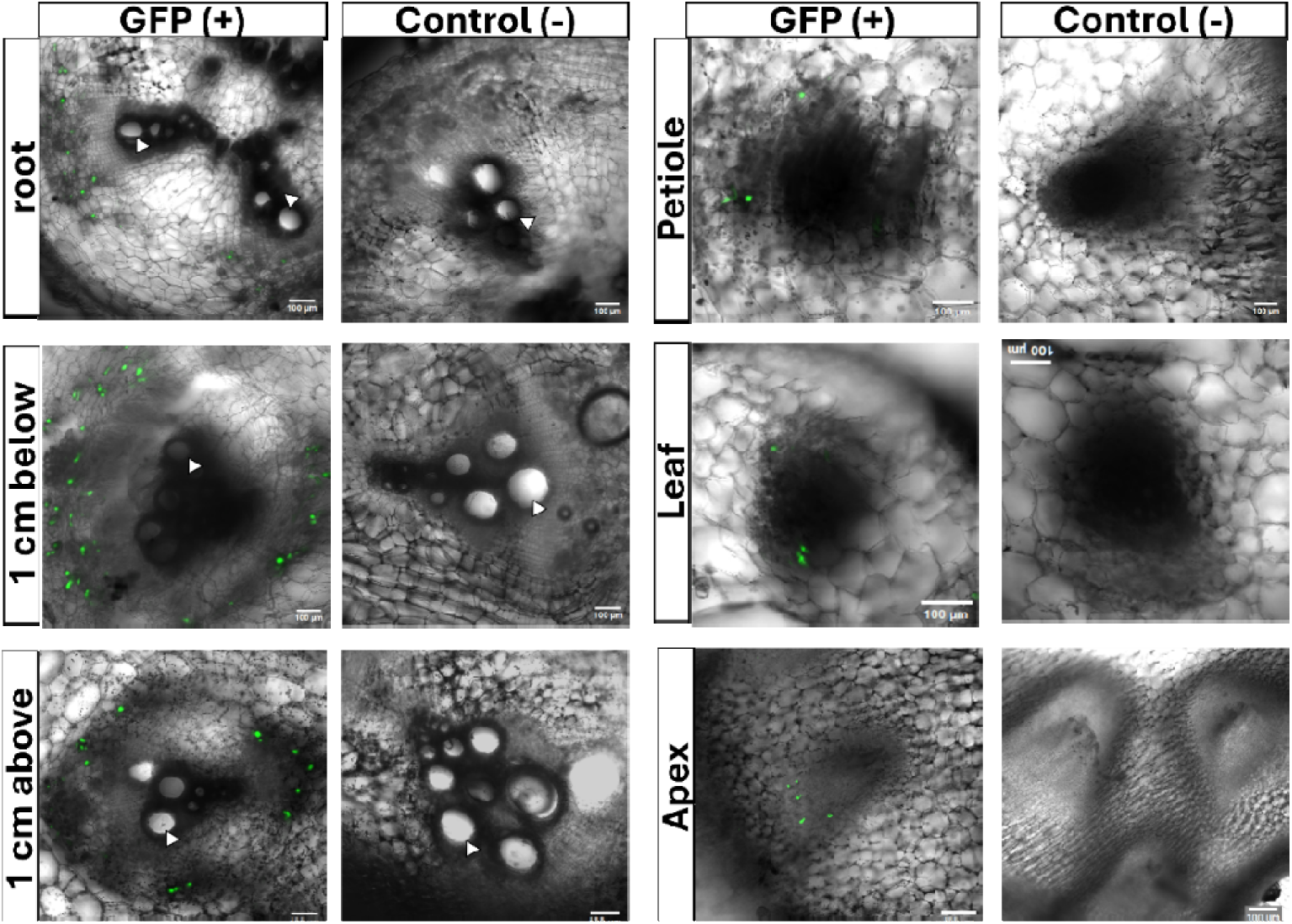
Confocal images show the localization of GFP-tagged *Serratia ureilytica* isolates across six tissue types. Representative images were selected for each tissue to illustrate bacterial localization within the anatomical context of the vascular bundle. Images were acquired at 10× magnification and are shown as merged brightfield and GFP channels. Brightness and contrast were adjusted on a per-image basis for visualization to facilitate detection of low-intensity GFP signals, where present; control images were displayed with intensity scaling matched to the most dimly fluorescent GFP-positive sample. Some images were cropped to highlight the anatomical region of interest. Post-acquisition adjustments were limited to display scaling and were not used for quantitative analysis. *white arrow indicates xylem

**Fig. 4.**
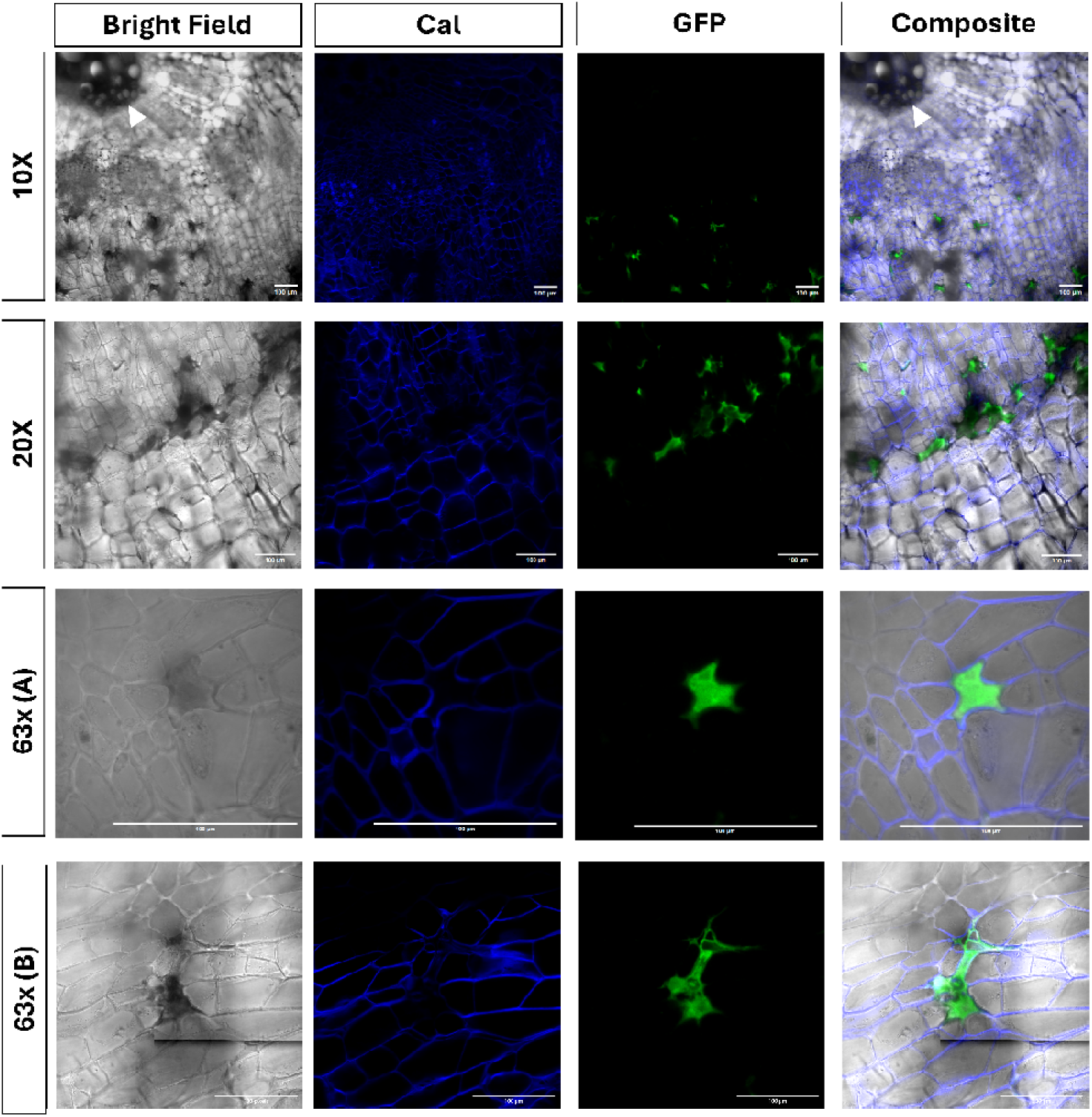
Confocal images of the same sample (1 cm below soil line) of *Cucurbita pepo* var. Delicata, acquired at 10×, 20×, and 63× magnifications to demonstrate symplastic and apoplastic (63X (B)) localization of GFP-tagged *Serratia ureilytica* relative to host cell walls. Tissue was collected 2 cm below the soil surface at 4 weeks post-inoculation and stained with calcofluor white immediately prior to imaging. From left to right, panels show brightfield, calcofluor white-stained cell walls (blue), GFP fluorescence (green), and merged composite images of all three channels. *white arrow in 10x indicates xylem

### Quantification of *S. ureilytica*

For the GFP P01 isolate, quantification results yielded plants in which *S. ureilytica* was isolated from one or more, but not all, sample locations, as well as plants where the pathogen was isolated from all sampling locations. Therefore, data analyses of bacterial colonization were performed using two datasets. First, analyses were performed on plants with *S. ureilytica* growth in at least one plant sampling location (**Fig 5**). A second analysis was performed comparing only plants with growth from all plant locations (**Fig. 6**). In the first analysis, results showed that at seven days post-inoculation (DPI), colonization by the GFP-tagged *S. ureilytica* P01 isolate was successful but more localized to the inoculation site (**Fig. 5**). There were high bacterial counts from the stem sections (1 cm below, 1 cm above, and 2 cm above the inoculation point), but no significant differences in log CFU/mL (**Fig. 5**). At 14 DPI, *S. ureilytica* was again isolated from stem sections, but the bacterium was also isolated from petioles. (**Fig. 5**). At 21 and 28 DPI, there was a higher detection rate from distal tissue, with stem sections at 1 and 2 cm above the inoculation point having higher log CFU/ml compared to the petiole, leaf, or apex tissue (**Fig. 5**). No *S. ureilytica* growth was seen in the mock-inoculated and uninoculated controls.

**Fig 5.**
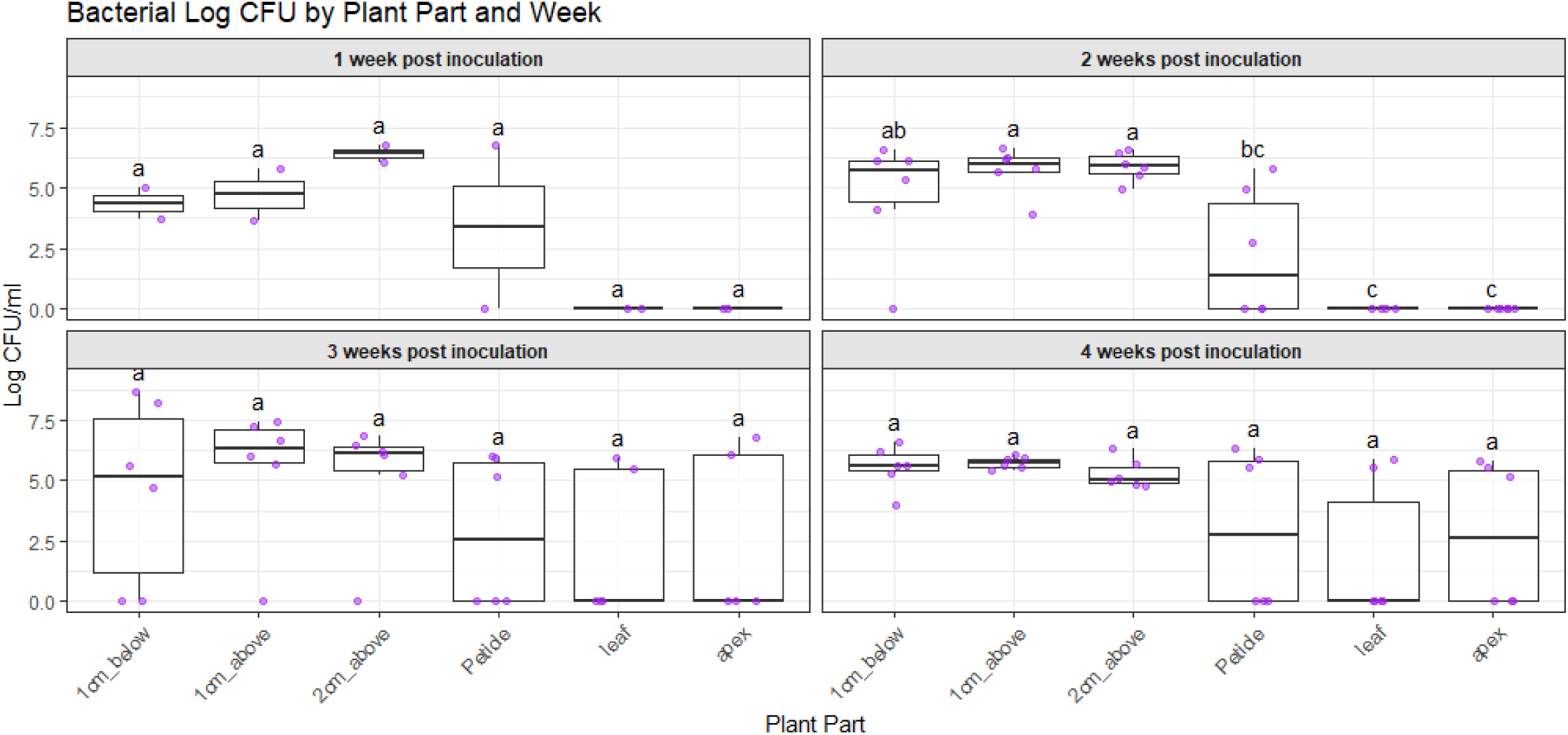
Log CFU/ml of *Serratia ureilytica* (CFU/ml) isolated from *Cucurbita pepo* plants inoculated at two weeks old and sampled weekly for four weeks. Each plant was sampled in one of six locations shown on the y-axis. Bars with the same letter are not significantly different from each other (P > 0.05) according to Tukey’s honestly significant difference. Error bars indicate standard error.

**Fig. 6.**
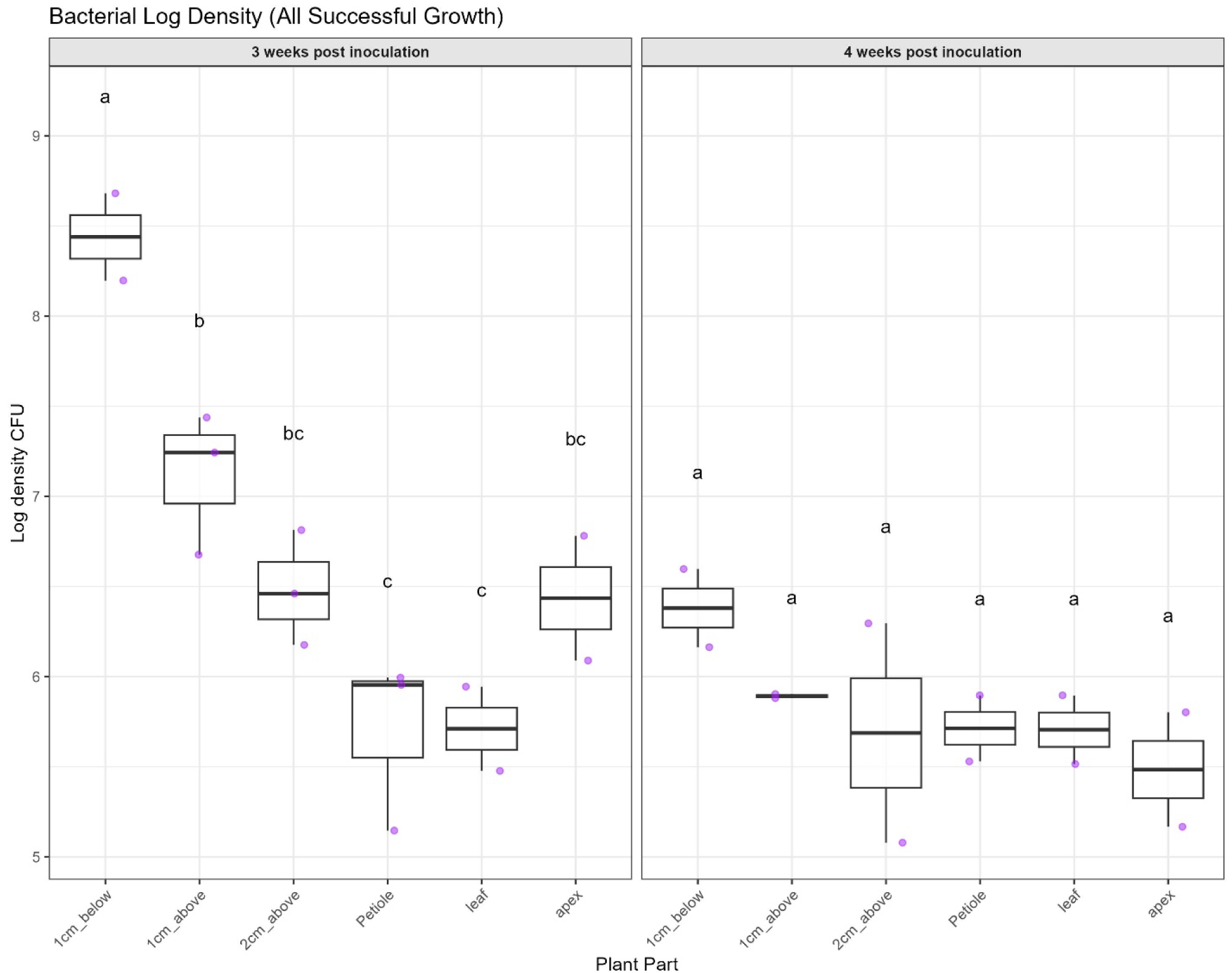
Log CFU/ml of *Serratia ureilytica* isolated from *Cucurbita pepo* plants inoculated at two weeks old, and sampled weekly for four weeks, including only those plants in which *S. ureilytica* colonies were detected in all six plant parts. Each plant was sampled in one of six locations shown on the y-axis. Bars with the same letter are not significantly different from each other (P > 0.05) according to Tukey’s honestly significant difference. Error bars indicate standard error.

Analysis limited to plants with successful growth in all parts showed that *S. ureilytica* colony counts were higher at the crown level than at other sampled locations (**Fig. 6**). At 21 DPI, *S. ureilytica* colony counts were significantly higher (log CFU/ml) in the three stem sections compared to all other plant tissues, with distal tissues having the lowest log CFU/ml (**Fig. 6**). At 28 DPI, bacterial distribution was not significantly different across sample locations, and the mean log CFU/ml showed a more even bacterial distribution across the plant with no significant differences among plant tissues (**Fig. 6**).

## Discussion

The present study aimed to understand the *in planta* localization and temporal distribution of *S. ureilytica,* the causal organism of CYVD. To achieve this, cucurbit plants (*C. pepo*) were inoculated at the cotyledon stage using a GFP-tagged *S. ureilytica* isolate. Dissection and confocal laser scanning microscopy revealed that *S. ureilytica* colonized specific cells at the interior and exterior peripheries of the vascular bundles of cucurbits, suggesting that bacterial colonization of *C. pepo* plants is restricted to phloem-associated tissues as previously hypothesized (Bruton et al. 1998a). This localization differs from *E. tracheiphila*, which is found in the xylem vessels of cucurbit plants, causing rapid water deprivation and wilt, in contrast to the slow symptom development caused by CYVD (Bruton et al. 1998a; Rojas et al. 2015). The presence of *S. ureilytica* was observed across time and location. We found strong evidence of symplastic localization, with putative GFP signal also localized to the apoplast.

Quantitative analysis of *S. ureilytica* showed that this bacterium moves within the plant over time. Results also suggest that the bacterium travels bidirectionally through the phloem sieve elements from the main stem to the root, up to the crown, and eventually throughout the plant. This is consistent with previous reports, including in vitro studies showing that Type I fimbriae present in *S. ureilytica* impedes this pathogen’s biofilm formation, thereby allowing a larger population of bacteria to travel through the plant’s phloem and colonize more tissue (Mphande et al.,2025a). Results from bacterial quantification in six plant tissues indicated that the pathogen was mainly found in stem tissue near the inoculation site for the first 14 days, then moved to more distal locations by 21 and 28 DPI, with titers that are not significantly different between sampling locations at 28 DPI. This might be explained by *S. ureilytica* fully colonizing phloem cells as it moves through the plant, which can also lead to increased symptom development, as higher populations of this bacterium are required to see CYVD symptoms (Mphande et al. 2025a).

Based on the results presented here, we hypothesize that *S. ureilytica* utilizes the sieve tubes of the phloem for transport, but colonizes other cells within the phloem. These findings are consistent with the leakage retrieval mechanism of phloem transport in which solutes in transport sieve tubes can "leak” out of sieve tubes, either into the apoplast or, in some conditions, through PD into companion cells (De Schepper et al., 2013).

Although we do not observe a GFP signal in sieve tube cells, bacterial presence in those cells may be transient and at concentrations too low to be detected. Fluorescence in our images is mainly symplastic, scattered throughout a subset of small to medium, dark, irregularly shaped cells, within the phloem-associated regions of the vascular bundle. We hypothesize that *S. ureilytica* could be colonizing one of two cell types: phloem parenchyma or companion cells. Companion cells are typically small and have dense cytoplasm, which can make them appear dark in brightfield images (Esau 1969). Companion cells also tend to be scattered throughout the phloem region of the vascular bundle, similar to what we see in our dissecting scope images.

However, companion cells are always directly associated with a sieve cell due to shared lineage, and it is not clear that our cells consistently show this phenotype. These cells may appear dark and irregularly shaped as a result of the infection, making them difficult to identify based on morphology alone. This localization differs from that of other known phloem colonizing bacteria, such as *Candidatus Liberibacter, asiaticus* in citrus, which is restricted to the sieve tube elements of the phloem (Ding et al. 2015; Tiwari 2025). Other examples include *Candidatus Arsenophonus phytopathogenicus* and *Candidatus Phytoplasma asteris,* which are also found in phloem sieve tube elements and are unculturable. This resistance to culturing may be explained by the specialization required for the bacteria to occupy sieve tubes, an environment with an extremely high nutrient concentration. This distinction may also explain why *S. ureilytica,* can be isolated on artificial media when other phloem colonizers cannot. These results contrast with previous studies that showed the presence of *S. ureilytica* only in phloem sieve tube elements (Bruton et al. 1998b).The fact that *S. ureilytica* is localized on phloem-associated cells raises the questions as to how both squash bug and cucumber beetles are able to deliver the bacterium into these cells. It is well known that the squash bug is able to feed from phloem tissue (Neal 1993).On the other hand, cucumber beetles use a stercorarian transmission, where they acquire the bacterium through feeding and deposit the infected frass into open chewing wounds when feeding (Mphande et al. 2025b). This infection process could involve using the polymer trapping loading system to deliver *S. ureilytica* from the leaves to the main stem which utilizes a concentration gradient to move sugars cell-to-cell via plasmodesmata into the phloem transport stream, using intermediary cells (Reidel et al. 2009; Turgeon 2017; Turgeon and Hepler 1989). Furthermore, previous research has shown that *Serratia* species that colonize insects proliferate at higher rates within the host when fed a high-sucrose diet in fruit flies (*Drosophila melanogaster*)(Darby et al. 2024). Given the high sucrose content in both companion cells and phloem parenchyma and in the apoplast compared to other plant tissues, this could drive the pathogen’s localization to these cells. Since sucrose is an important carbon source for nitrogen assimilation in plants (Huang et al. 2022), hijacking sucrose may explain CYVD symptoms such as stunting and yellowing of older leaves (Besler 2014; Bruton et al. 1998a; Pair et al. 2004).

In both the imaging and quantification experiments, we consistently observe greater bacterial abundance in the lower tissues of the main stem and root. This is consistent with the dynamics of bulk flow in the phloem. Bulk flow, or the movement of sap through the phloem, is powered by the pressure gradient, driven by osmotic forces due to varying concentrations of solute in the sap (high concentrations in mature leaves where photosynthesis occurs and low concentrations in storage tissues like roots) (Fellows and Geiger 1974). The direction of flow in any given sieve tube changes throughout development, with architectural modifications such as the initiation and maturation of new leaves and the growth of the root system, but during vegetative growth stages apical-to-basal flow dominates. Furthermore, it is established that some phloem-occupying pathogens can modify metabolic activity in colonized cells, thereby altering transport direction and increasing nutrient delivery to infected organs (Malinowski et al., 2024). These dynamics are consistent with the localization of *S. ureilytica* observed in our plants, as distal tissues are more likely to be occupied at higher densities at later time points, with the majority of bacterial growth occurring in the main stem and root. This work provides novel evidence of *S. ureilytica* localization in cucurbit plants and advances our knowledge of this emergent disease in the US. Building on these findings, microscopy methods, including histology and immunolocalization can be used to confidently establish cell-type localization within the phloem. Given *S. ureilytica’s* preference to colonize phloem companion cells or phloem parenchyma cells and move systemically throughout the plant, further research should prioritize elucidating the molecular mechanisms driving this localization. Future studies could investigate environmental cues, such as sucrose gradients, to determine if they act as chemoattractants for the bacterium within the phloem. Moreover, further investigation into the interaction between *S. ureilytica* and cucurbit plasmodesmata could clarify if the bacterium actively manipulates symplastic transport pathways or relies on passive movement.

## Acknowledgements

We thank Elizabeth Little and Nicole Gerardo for sharing the GFP *S. ureilytica* isolate used in this study.

---

**Supplementary Fig. 1:**
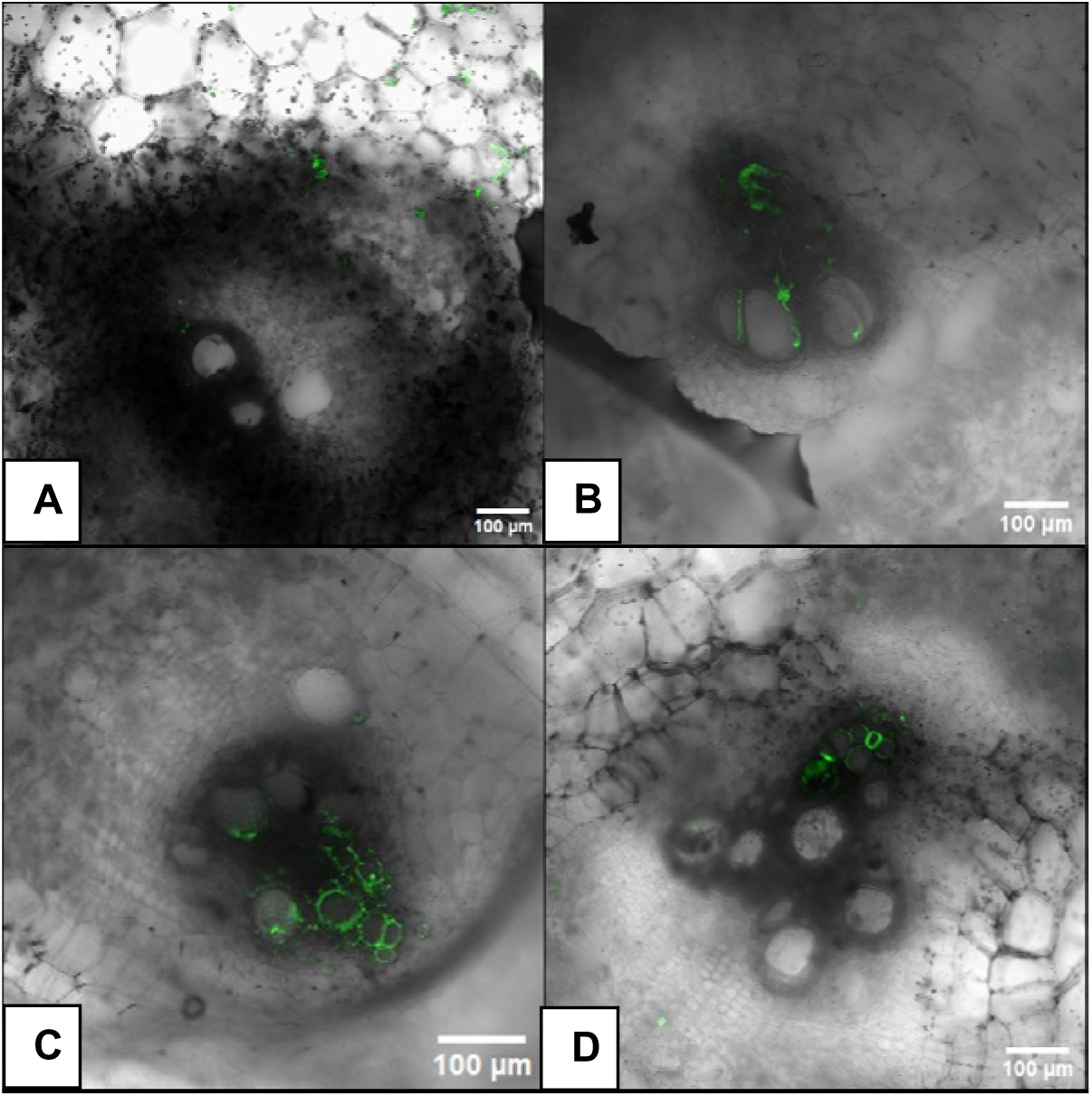
autofluorescence in control images using confocal microscopy at 10x. Fig. A. Autofluorescence in the chloroplast at 20X. Fig. B-D Autofluorescence in xylem tissue. GFP Gain A:985 B: 559. C: 718. D: 705

